# A map of constrained coding regions in the human genome

**DOI:** 10.1101/220814

**Authors:** James M. Havrilla, Brent S. Pedersen, Ryan M. Layer, Aaron R. Quinlan

## Abstract

Deep catalogs of genetic variation collected from many thousands of humans enable the detection of intraspecies constraint by revealing coding regions with a scarcity of variation. While existing techniques summarize constraint for entire genes, single metrics cannot capture the fine-scale variability in constraint within each protein-coding gene. To provide greater resolution, we have created a detailed map of constrained coding regions (CCRs) in the human genome by leveraging coding variation observed among 123,136 humans from the Genome Aggregation Database (gnomAD). The most constrained coding regions in our map are enriched for both pathogenic variants in ClinVar and de novo mutations underlying developmental disorders. CCRs also reveal protein domain families under high constraint, suggest unannotated or incomplete protein domains, and facilitate the prioritization of previously unseen variation in studies of disease. Finally, a subset of CCRs with the highest constraint likely exist within genes that cause yet unobserved human phenotypes owing to strong purifying selection.

## INTRODUCTION

During World War II, Abraham Wald and the Statistical Research Group optimized the placement of scarce metal reinforcements on Allied planes, based on the patterns of bullet holes observed over the course of hundreds of sorties. Wald famously invoked the principles of survival bias to intuit that armor should be placed where bullet damage was *unobserved*, since the observed damage came solely from planes that *returned* from their missions. Wald reasoned that planes that had been shot down likely took on critical damage in such locations^1^.

Employing similar logic, we sought to identify localized, highly constrained coding regions in the human genome. We were motivated by the idea that the absence of genetic variation in coding regions (e.g., one or more exons or portions thereof) ascertained from large human cohorts implies strong purifying selection owing to essential function or disease pathology. An intuitive approach to identifying intraspecies genetic constraint on human coding genes is to identify gene sequences that harbor significantly less genetic variation than expected. For example, Petrovski et al. used genetic variation observed among 6,515 exomes in the NHLBI Exome Sequencing Project dataset^2^ to develop the Residual Variation Intolerance Score (RVIS)^3^, which ranked genes by their intolerance to “protein-changing” (i.e., missense or loss-of-function and coding) variation. Similarly, Lek et al. recently integrated variation observed among 60,706 exomes in the Exome Aggregation Consortium (ExAC)^4^ to estimate each gene's probability of loss-of-function intolerance (pLI), with genes having the highest pLI harboring significantly less loss-of-function (LoF) variation than predicted^5^.

While existing gene-wide metrics of constraint are effective for disease variant interpretation, single metrics cannot capture variability in regional constraint that exists within protein-coding genes. Constraint variability is expected given that some regions encode conserved domains^6–10^ critical to protein structure or function, while others encode polypeptides that are more tolerant to perturbation. Therefore, while useful, single gene-wide metrics such as pLI are susceptible to both over- (Figure 1A) and underestimating (Figure 1B) local constraint within genes exhibiting finer-scale variation in constraint. Consequently, they are incapable of highlighting the subset of critical regions within each gene that are under the greatest selective pressure (Figure 1, regions highlighted in red). This manuscript presents a detailed map of constrained coding regions (CCRs) in human genes. We demonstrate that the most constrained regions recover known disease loci, empower variant prioritization, and illuminate new genes that may underlie previously unknown disease phenotypes. This map is shared openly and will only improve as ever larger catalogs of genetic variation are created.

**Figure 1.**
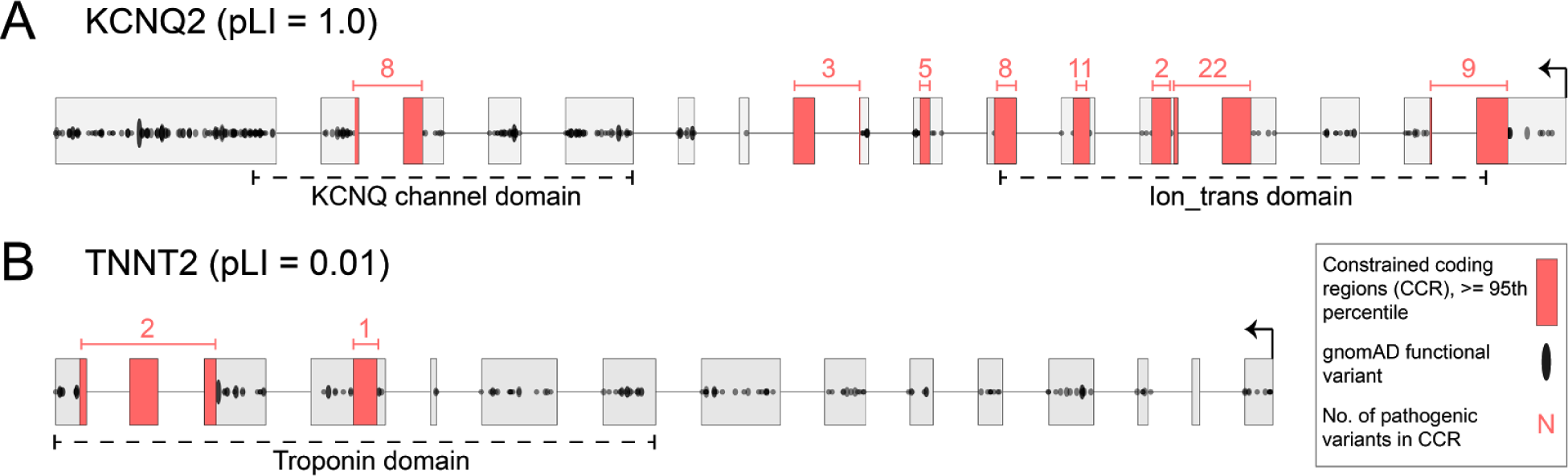
Gene-wide summary measures of constraint are prone to over- and understating the constraint that exists within specific regions of protein coding genes. (**A**) KCNQ2 has the highest possible pLI score of 1.0, yet there are entire exons (e.g., the leftmost exon) with many protein-changing variants indicating they are under minimal constraint. Highly constrained (i.e., in the 95th percentile or higher as described in text) coding regions highlighted in red are completely devoid of protein-changing variation in gnomAD. (**B**) In contrast, TNNT2, which regulates muscle contraction and has been implicated in familial hypertrophic cardiomyopathy^11^, has a very low pLI of 0.01. However, there are focal regions lacking protein-changing variation, indicating a high degree of local constraint. Numbers above each constrained coding region (CCR) reflect the number of ClinVar pathogenic variants in each CCR, and illustrate that CCRs often coincide with known disease loci.

## RESULTS

### Constructing a map of constrained coding regions (CCRs)

Hypothesizing that coding regions under extreme purifying selection should be devoid of protein-changing variation in healthy individuals, we have created a high-resolution map of constrained coding regions (CCRs) in the human genome. The gnomAD database (v.2.0.1) reports 4,798,242 missense or loss-of-function variants among 123,136 human exomes, yielding an average of 1 variant every ˜7 coding base pairs. Given this null expectation of high protein-changing variant density, we sought to identify the exceptions to the rule: that is, coding regions having a greater than expected distance between protein-changing variants owing to constraint on the interstitial coding region (Figure 1, regions highlighted in red). Simply stated, CCRs having no protein-changing variation over the largest stretch of coding sequence (weighted by sequencing depth) are assigned the highest percentiles and are inferred to be under the highest constraint in the human genome.

Our CCR map is charted by first measuring the exonic (ignoring introns) distance between each pair of protein-changing gnomAD variants found in autosomal genes. Each region's “length” is then weighted by the fraction of gnomAD samples having at least 10X sequence coverage in the region. This correction prevents the false identification of constraint arising simply because poor sequencing coverage reduced the power to detect variation. Similarly, we exclude coding regions that lie in segmental duplications or high identity (>=90%) self-chain repeats^12^ to avoid the confounding effects of mismapping short reads among paralogous coding regions^13^. After these exclusions, we are able to measure localized constraint for 88% of the protein coding exome. For each region, we adjust for the CpG dinucleotide density as an independent measure of the potential mutability of the coding region.^14^ While other models^5,15,16^ of local mutability have been developed, the primary predictor of these studies and others^17,18^ is the presence of CpG dinucleotides. We fit a linear model of the region's CpG density (the independent variable) versus its weighted length (the dependent variable). The greater the difference between the observed and expected weighted length for regions with similar CpG density, the higher the constraint. Finally, each coding region is assigned a residual percentile that reflects the degree of constraint, where higher percentiles reflect greater predicted constraint (**Methods**). A gene's coding sequence length is weakly correlated (Pearson r=0.28, Supplemental Figure 1) with the maximum CCR percentile observed in the gene, indicating, as expected, that a gene's size is a minor contributor to the probability of observing a highly constrained region. Furthermore, the median exonic length of CCRs in the 95th (52 bp) and 99th (94 bp) is far greater than the 7 bp expected distance between protein-changing variants (Supplementary Figure 2).

### Constrained coding regions are enriched in disease-causing loci

To evaluate the relationship between CCRs and loci known to be under genetic constraint, we measured the enrichment of pathogenic ClinVar variants (**Methods**) versus ClinVar benign variants across CCR percentiles. As expected, pathogenic variants from all disease types are significantly enriched in the 95th CCR percentile and above (OR=161.8; 95% CI=[40.4 - 647.5]), and depleted (OR=0.019; 95% CI=[0.015 - 0.023]) in the least constrained coding regions (Figure 2A, light green bars). Not surprisingly, given that CCRs identify coding regions that lack any protein-changing variation in the gnomAD database, pathogenic variants for autosomal dominant disorders are even more enriched in the 95th CCR percentile and above than other disorders (OR=225.4; 95% CI=[56.3 - 902.7]) (Figure 2A, dark green bars).

The most constrained regions are restricted to a small fraction of genes. Of the 17,693 Ensembl^19^ genes in the CCR model, merely 8.0% and 3.9% of genes have at least one CCR in the 99th and 99.5% percentile or higher, respectively (Figure 2B). Genes exhibiting multiple highly (99th percentile or higher) constrained regions include many known to be involved in developmental delay, seizure disorders, and congenital heart defects, including KCNQ2, KCNQ5, SCN1A, SCN5A, multiple calcium voltage-gated channel subunits (e.g., CACNA1A, CACNA1B, CACNA1C, etc.), and GRIN2A (**Supplementary Table 1**). In addition, nine chromodomain helicase DNA-binding (CHD) genes and the actin-dependent chromatin regulator subunits SMARCA2, SMARCA4, and SMARCA5 contain multiple 99th percentile CCRs. Such constraint likely reflects their role in chromatin remodeling, development, and severe disorders.^20, 21^ Finally, while highly constrained regions often contain one or more known pathogenic variants, more than 2000 CCRs in the 99th percentile do not overlap a known pathogenic variant (Figure 2C). We hypothesize that this is likely owing to the fact that these regions are under extreme purifying selection which prevents the observation of a pathogenic variant among patients studied to date. There is support for this hypothesis; genes predicted to be essential, despite not having a known disease association,^22^ are significantly enriched relative to nonessential genes in the set of 1,415 genes with at least one 99th percentile CCR (one-tailed Fisher's exact test, p=8.6e-67, OR=3.73). Furthermore, genes lying in loci with low haplotype diversity are enriched for essential function^23,24^ and are similarly prevalent in genes having a 99th percentile CCR. This enrichment is significant when compared to genes with higher haplotype diversity (one-tailed Fisher's exact test, p=5.6e-5, OR=4.4). Taken together, these findings suggest that genes lacking a disease association yet harboring one or more 99th percentile CCRs are under strong purifying selection owing to extreme fitness consequences when mutated.

**Figure 2.**
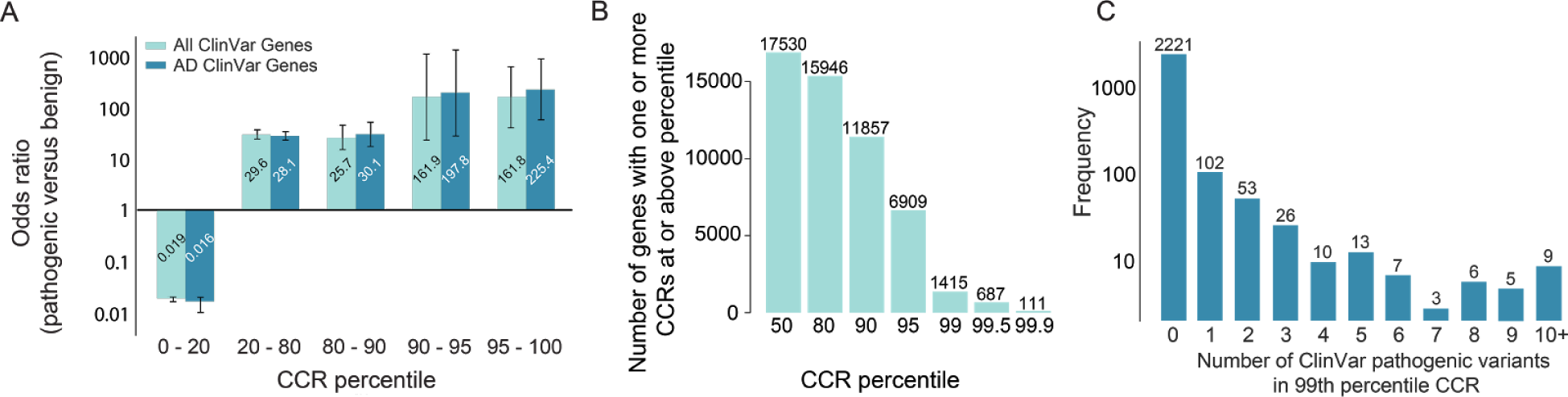
The most constrained coding regions are enriched for pathogenic variants and restricted to a small subset of genes. (**A**) Odds ratio enrichment for ClinVar pathogenic variants versus benign variants for different CCR percentile bins across all Clinvar genes (light green), as well as genes that underlie autosomal dominant (AD) diseases (dark green). (**B**) Histogram of the number of genes with at least one CCR greater than or equal to different percentile thresholds. (**C**) Histogram of the number of 99th percentile CCRs with 0 to 10 or more overlapping ClinVar pathogenic variants. CCRs in the 99th percentile that harbor no known pathogenic variants may reflect regions under extreme purifying selection.

### Comparing intraspecies and interspecies constraint

Given that most human genes are conserved among vertebrates, it is logical to expect that intraspecies constraint would be correlated with interspecies conservation, and that the most constrained coding regions would lie within conserved protein domains. To explore the relationship between intraspecies constraint and interspecies conservation, we compared CCRs to mammalian conservation measured by GERP++^25^ (Figure 3A). Overall, constraint is weakly correlated (Pearson r=0.002 overall, r=0.30 for CCRs of length 20 and higher) with conservation, illustrating that intraspecies constraint complements, and is not merely a subset of, conservation measures. As expected, the majority (98.2%) of the most constrained CCRs (95th percentile and above) have mean GERP++ scores that suggest conservation in vertebrates (>0.7 mean GERP score). However, 399 (Figure 3A, in dotted box) CCRs in 360 distinct genes are weakly conserved and suggest that some of these regions may represent recent constraint within the primate or human lineage. For example, CDKN1C contains a 98.3 percentile CCR (96 bp without variation in gnomAD) that coincides with a ClinVar variant known to be pathogenic for Beckwith-Wiedemann Syndrome^26,27^. CDKN1C is imprinted with preferential expression of the maternal allele^28^, suggesting that monoallelic expression may, in part, underlie the degree of observed constraint as the expression of only one allele opens greater risk for a dominant phenotype. Furthermore, our model includes 30 of the 42 imprinted genes reported by Baran et al. using data from the Genotype-Tissue Expression (GTEx) project^29^. Sixteen of 30 (53%) imprinted genes harbor at least one CCR in the 95th percentile or higher: GRB8, IGF2, KCNQ1, KIF25, MAGEL2, MAGI2, MEST, NAP1L5, NTM, PEG10, PEG3, PLAGL1, SNRPN, SYCE1, UBE3A, and ZDBF2. This reflects a 1.35 fold enrichment over the 39% (6909 of 17693) of all genes in the CCR model having a CCR in at least the 95th percentile. Other genes harboring similarly dichotomous constraint and conservation measures include four members of the Fanconi anemia pathway (FAN1, SLX4/FANCP, BOD1L1, and ERCC5), as well as an overrepresentation (p=7.9e-06, **Methods**) of genes involved in the complement cascade of the innate immune system (**Supplementary Table 2**).

Motivated by prior analyses^30^, we then explored the landscape of constraint in Pfam^31^ domains, given that protein domains are conserved owing to their structural or functional role in proteins (**Supplementary Document 1**). While constraint is often uniformly distributed over many protein domains (Figure 3B), several families are enriched for high constraint, likely owing to their critical function in proteins that contain them (Figure 3C). Constraint within ion transport domains is expected given their role in regulating the critical specificity of ion transport and the fact that mutations in these domains cause autosomal dominant encephalopathies^32^, neuropathies^33^ and cardiomyopathies^34^. Furthermore, homeobox domains bind DNA and are involved in cellular differentiation and maintaining pluripotency.^35^ Helicase superfamily C-terminal domains catalyze DNA unwinding and are implicated in Alpha-thalassemia^36^ and mental retardation^37^. Moreover, PHD finger domains are found in many chromatin remodeling proteins, which, when perturbed, lead to various disorders^38–40^. Finally, the eIF-5a domain is solely found in the two EIF5A translation initiation factors. These are the only human proteins that utilize the rare amino acid, hypusine. Strikingly, a 99.47 percentile CCR coincides with the hypusine in the primary isoform of EIF5A. Knockout of either the EIF5A gene or the deoxyhypusine synthase gene, whose product is necessary to create the hypusine amino acid for EIF5A, causes embryonic lethality in mice^41^.

It is notable that 43% (24,431 of 57,408) and 31% (759 of 2,454) of 90th and 99th percentile CCRs do not coincide with an annotated Pfam domain. While CCRs that are proximal to annotated Pfam domains likely reflect truncated annotations owing to reduced homology or the consequence of homology searches driven by local alignment, distal CCRs may represent coding regions of previously uncharacterized functional or structural importance.

**Figure 3.**
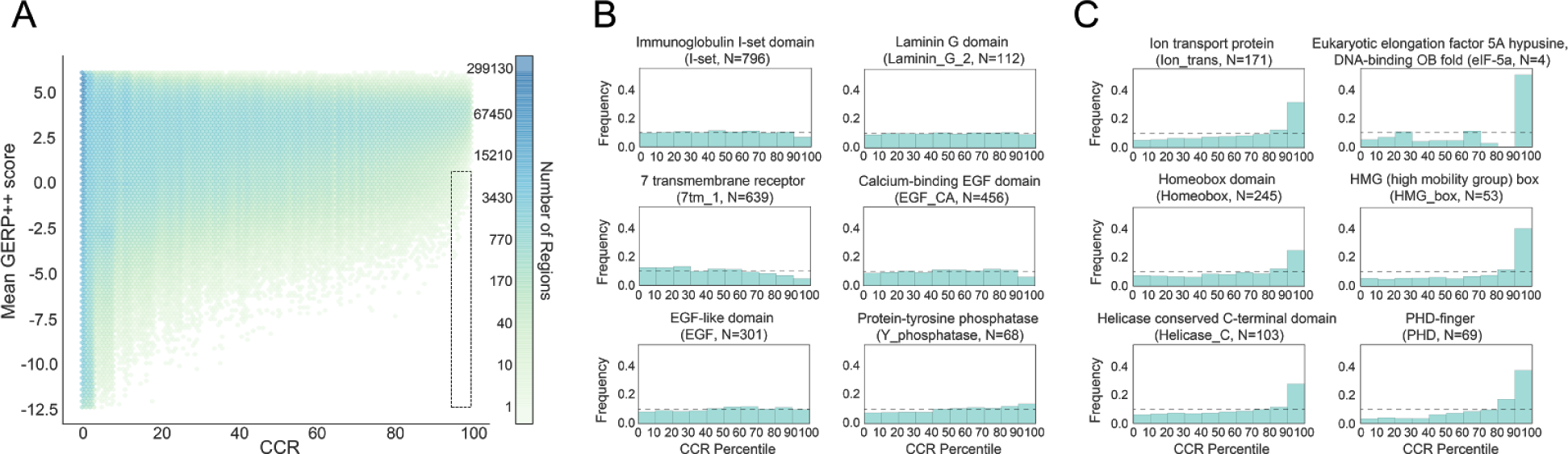
(**A**) A comparison of intraspecies constraint (CCR) versus interspecies conservation, as measured by the mean GERP++ score in each CCR. Regions in the dashed box reflect intraspecies constraint not revealed by interspecies conservation: that is, they have less than 0.7 GERP++ score, and 95th percentile or greater CCR score. (**B**) Example Pfam domain families for which constraint is nearly uniformly distributed among instances of the domain. (**C**) Representative Pfam domain families exhibiting enrichment for higher levels of intraspecies constraint across the whole exome.

### Comparing CCRs to other models of regional constraint

Samocha et al. recently described an approach to identify regions of protein coding genes that exhibit “missense depletion”: that is, regions where far less than expected missense variation is observed in the Exome Aggregation Consortium (ExAC v1) catalog of 60,706 exomes^42^. While the motivation is similar to our model of regional constraint, the missense depletion approach partitions solely 15.1% of transcripts into distinct missense depletion regions. That is, for 85% of transcripts, the entire transcript is assigned a single, summary constraint measure, and only 5.5% of transcripts are partitioned into three or more distinct regions of missense depletion. The missense depletion approach also chooses a single representative transcript for each gene, thus coding exons exclusive to other isoforms are not modeled. Since CCRs measure constraint variability along the entire gene, they provide a more detailed map of the spectrum of constraint and identify additional highly constrained coding regions. As a result, 1,874 of the top 1% most constrained CCRs would be classified as either “unconstrained” or “moderately constrained” by the missense depletion threshold (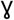 > 0.4) (Figure 4). These regions lie within 800 distinct genes, many of which have known associations with autosomal dominant disease (**Supplementary Table 3**). Therefore, the two models of regional constraint provide complementary information, since CCRs provide a detailed constraint architecture for 88% of the exome, whereas missense depletion delineates regional constraint for 15% of the protein coding transcripts.

**Figure 4.**
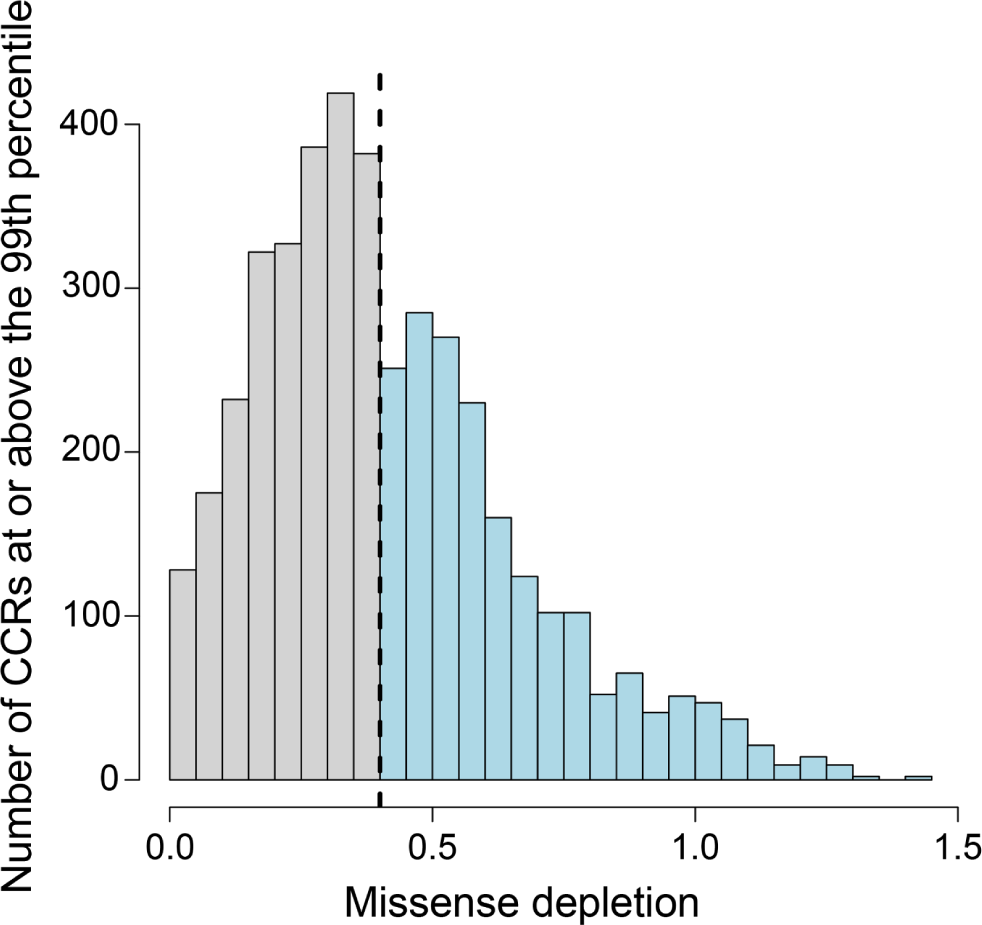
Constrained coding regions provide greater detail in local coding constraint than missense depletion regions. The dashed line reflects the missense depletion threshold (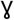 > 0.4) below which Samocha et al. define regional constraint. Light blue bars above this threshold reflect CCRs at or above the 99th percentile that would not be deemed as constrained by the missense depletion metric. Grey bars reflect CCRs that coincide with regions deemed to be under constraint by missense depletion.

### Using constrained coding regions to prioritize variants in disease studies

Given the explosive human population growth over the last two millennia^43^ and the resulting excess of very rare genetic variation in the human genome, a natural question is the degree to which constraint measured from >120,000 exomes is sufficient to empower the prioritization of variation observed in newly-sequenced individuals with disease. Since the highest CCRs are, by definition, devoid of protein-changing variants observed even as a heterozygote in a single individual from gnomAD, we should expect them to harbor mutations in de novo dominant disorders. We tested this hypothesis by comparing the enrichment of 5,113 de novo missense mutations (DNMs) in 5,620 neurodevelopmental disorder probands^44–49^ (“pathogenic” variants) versus 1,269 missense DNMs from 2,078 unaffected siblings of autism spectrum disorder probands^50,51^ (**Table S8** from Samocha et al.^42^). These mutations were assumed to be benign and serve as control DNMs. Strikingly, we observe a 7.1 fold enrichment of DNMs from neurodevelopmental disorder cases in the most constrained CCRs and a 0.25 fold depletion of DNMs in cases in the least constrained CCRs (Figure 5A).

We then compared the performance of CCRs for variant prioritization to GERP, CADD^52^, REVEL^53^ and MPC^42,53^. When tested on the same ClinVar pathogenic and benign variants described above (**Methods**), the CCR model yields the highest area under curve (AUC, 0.94) for the receiver operating characteristic (ROC) curve analysis (Figure 5B) and the highest Youden J statistic^54^. ClinVar represents a highly curated truth set that is an optimistic proxy for the variant prioritization challenges inherent to actual studies of rare disease. Therefore, we also tested the performance of each metric using the 5,113 putative pathogenic and 1,269 putative benign DNMs from Samocha et al. While not all DNMs from the neurodevelopmental disorder probands are truly pathogenic and not all DNMs in unaffected siblings are necessarily benign, we expected to see enrichment for truly pathogenic variants in the de novo variants from samples with developmental disorders relative to controls. While the performance of all methods is reduced on this more difficult evaluation set, the CCR model yields the highest ROC AUC (0.73) among tested methods. This reflects its additional power to recover coding regions under true constraint and score variants not found solely in ClinVar (Figure 5B).

**Figure 5.**
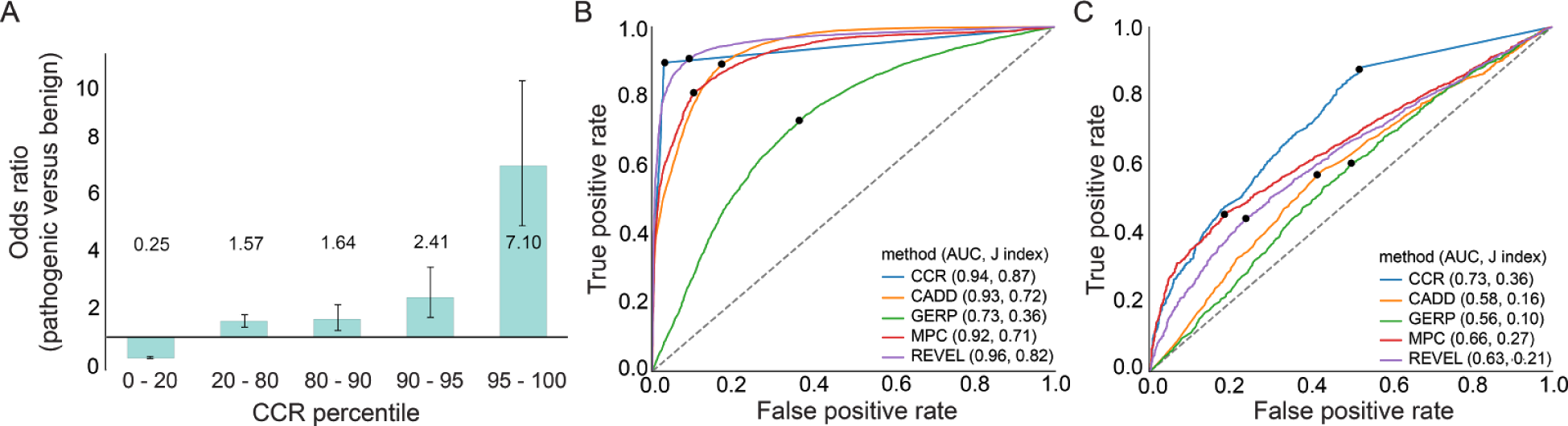
Evaluation of *de novo* variation from a cohort with severe developmental delay, intellectual disability, and epileptic encephalopathy versus *de novo* variation from unaffected siblings of autism probands. (**A**) Enrichment of pathogenic de novo mutations in the most constrained CCRs. (**B**) ROC curve for CCR versus other metrics for the ClinVar evaluation set. True positives are pathogenic variants and likely pathogenic variants from ClinVar. True negatives are variants labeled as benign from ClinVar. (**C**) ROC curve for developmental disorder *de novo* variant evaluation set. The true positives are missense-only *de novo* variants from patients with developmental disorders. The true negatives are missense *de novo* variants from unaffected siblings of autism patients. The dots in (**B**) and (**C**) indicate the score cutoff with the maximal Youden J statistic for each tool. Values in parenthesis indicate the AUC and the maximal J, respectively.

### Estimating the rate of false positive discovery of coding constraint

Our current model of regional coding constraint is based upon variation observed from 123,136 exomes. However, Zou et al. estimate^55^ that even 500,000 individuals will be insufficient to catalog the majority of protein-changing variants in the human population. Yet if predicted regions of constraint are truly under strong purifying selection, they should remain largely free of protein-changing variation, even as genetic variation is collected from much larger cohorts of healthy individuals. To test the predictive power of the current model, we compared CCRs to DNMs observed in both neurodevelopmental disorder probands and unaffected siblings of autism probands. We assumed that DNMs from neurodevelopmental disorder probands represent true positives and DNMs from unaffected siblings represent true negatives, and thus false positives when they lie in regions of highest constraint. We measured the false discovery rate of each CCR in the 90th, 95th, and 99th percentiles (Table 1, **Methods**). Merely 2.8% of CCRs in the 99th percentile and higher coincide with a DNM from an unaffected sibling (FDR), and only 0.6% of these ostensibly benign DNMs lie within a 99th percentile or higher CCR. This suggests that while many more genomes are necessary to reveal all variation in the human genome, our model illuminates coding regions under true constraint at a low false discovery rate. Furthermore, a fundamental strength of our approach is that the resolution of predicted constraint will improve as variation from ever-larger cohorts of healthy individuals is integrated in future versions.

**Table 1.**
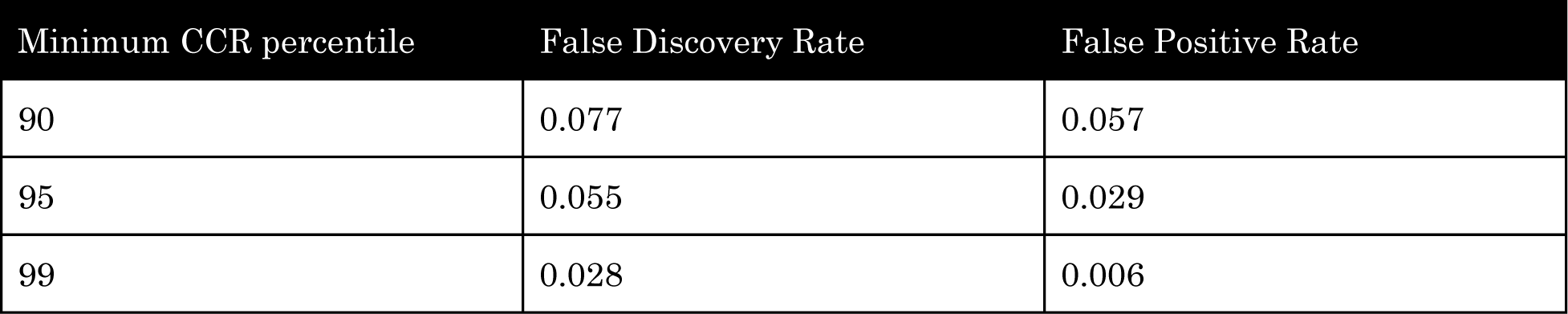
Estimated false discovery rate (FDR) and false positive rate (FPR) for the 90th, 95th, and 99th percentiles. De novo mutations from neurodevelopmental disorder probands are treated as true positives. De novo mutations from unaffected siblings of autism probands represent true negatives (TN), and when overlapping a CCR, are treated as false positives (FP). FDR is calculated as FP/(FP+TP) and FPR as FP/(FP+TN).

## DISCUSSION

Deep sampling of human variation provides a richly textured “topographical map” of constraint within protein coding genes. The map of constrained coding regions we have created reveals the broadest “valleys”; that is, local coding regions within genes that lack protein-changing variation from a sample of 246,272 human chromosomes. Our hypothesis posits that such regions are depleted for protein-changing variation because mutations therein have strong selective pressures against them. Supporting this hypothesis, we have shown that CCRs are enriched for disease-causing variants, especially in autosomal dominant Mendelian disorders. Furthermore, protein domains with critical function (e.g., ion transport, DNA binding, and chromatin remodeling) are enriched for the highest local constraint. These observations demonstrate the utility of CCRs for prioritizing variants in studies of rare human disease. While correlated, local coding constraint complements phylogenetic conservation measures. Therefore, building upon the work of Samocha et al.^42^, we argue that future improvements in variant prioritization will arise by combining models of local coding constraint with single-nucleotide metrics that incorporate complementary information such phylogenetic conservation, amino acid substitution scores, and 3D protein structure.

While we have demonstrated that highly constrained coding regions recover variants known to underlie human disease, we acknowledge that our approach is conservative. That is, by requiring the complete absence of protein-changing variation within a CCR, we are prone to false negatives in larger constrained regions where variation is extremely sparse yet not completely absent in healthy individuals. However, we sought to improve upon the resolution of existing gene-wide constraint measures and to minimize false positives by strictly identifying regions with the highest constraint within each gene. Consequently, CCRs have the greatest power for revealing regions of constraint under autosomal dominant disease models. Another important caveat of our model is that 55% (76,266 of 138,632) of the individuals sequenced in the gnomAD cohort are of European ancestry. Consequently, our current model of local coding constraint has lesser predictive power for non-European cohorts. Finally, our current CCR map excludes sex chromosomes owing to the reduced power to measure rare variation among a cohort comprised of males and females. We envision future extensions of our work that model this reduced power to provide maps of constraint in the X and Y chromosomes.

Perhaps the most useful outcome of a detailed map of coding constraint is the ability to reveal critical regions in genes that have not yet been linked to human disease phenotypes. We have shown that the most constrained regions are enriched for disease-causing variants (Figure 1A). However, nearly 91% of genes harboring at least one CCR in the 99th percentile or higher have no disease association in ClinVar. We hypothesize that some of these regions exhibit such extreme constraint because mutations therein are either incompatible with life or lead to extreme developmental disorders. Looking forward, investigating the phenotypic effects of disrupting these regions provides the opportunity to reveal new coding regions that underlie disease phenotypes and are vital to human function.

## DATA AVAILABILITY

Website: https://github.com/quinlan-lab/ccrhtml

Browser : https://rebrand.ly/ccrregions

BED file: https://s3.us-east-2.amazonaws.com/ccrs/ccrs/ccrs.v1.20171112.bed.gz

## ACKNOWLEDGEMENTS

We acknowledge William Pearson, Cedric Feschotte, Jon Seger, Gabor Marth, Nels Elde, and Stephanie Kravitz for insightful discussions that motivated some of the analyses presented in this manuscript. We also thank the investigators that contributed to the Genome Aggregation Database for openly sharing the genetic variation datasets that facilitated our research.

## AUTHOR CONTRIBUTIONS

ARQ conceived the research question and organized the study. JMH led the research and analysis. JMH, BSP, RLM, and ARQ designed the coding constraint region model and contributed to the analyses. JMH and ARQ wrote the manuscript.

## METHODS

### CCR model construction

The map of constrained coding regions (CCR) is constructed from the catalog of genetic variation observed among 123,136 exomes in gnomAD: (https://storage.googleapis.com/gnomad-public/release/2.0.1/vcf/exomes/gnomad.exomes.r2.0.1.sites.vcf.gz). We applied vt^56^ variant normalization and decomposition to the gnomAD VCF, followed by annotation with VEP^57^ (version 81 using Ensembl version 75 transcripts). The CCR model uses solely variants that VEP predicts to be “protein-changing”, which we define as any variant having the following Sequence Ontology terms for at least one Ensembl transcript: 'missense_variant', 'stop_gained', 'stop_lost', 'start_lost', 'frameshift_variant', 'initiator_codon_variant', 'rare_amino_acid_variant', 'protein_altering_variant', 'inframe_insertion', 'inframe_deletion', and ‘splice_donor_variant’ or ‘splice_acceptor_variant’ when paired with ‘coding_sequence_variant’. In addition, the variants must have a filter value of “PASS”, “SEGDUP”, or “LCR”. The rationale behind including “LCR” and “SEGDUP” labeled variants is that we already account for segmental duplications and self-chains in our model. In an effort to avoid excluding any real variants in gnomAD, we include variants with such filters and let our annotations of segmental duplications and self-chains exclude variants.

Coding exons from all protein coding transcripts in ENSEMBL^19^ version 75 were “flattened” into a single, combined model of coding sequence for each gene. Constraint “regions” are defined by measuring the exonic nucleotide distance between each pair of protein-changing variants. Therefore, constraint regions can exist within a single exon or span multiple exons. In order to prevent false identification of constraint that could arise solely because of reduced power to detect genetic variation, the length of each region is weighted by the fraction of individuals in gnomAD having at least 10x coverage at each bp. For example, if a region is 100 bp long and at each bp 90% of individuals have 10x coverage, the resulting weighted distance would be 90. Additionally, if the coverage falls below 50% of gnomAD individuals having at least 10x, the region is immediately broken and a new region is not started until the coverage exceeds 50% of individuals at 10X coverage. Finally, coding regions that overlap either segmental duplications or self-chain alignments with at least 90% identity are removed from our model. The rationale is that we cannot trust variant patterns in these regions owing to known artifacts that may arise when aligning short sequencing reads to paralogous genome segments.

For all remaining constraint regions, we compute the region's CpG density as a proxy for the region's mutability owing to spontaneous deamination of methylated cytosines. We then create a linear regression of the weighted length (dependent variable) versus CpG density (independent variable) for all regions. Each region's degree of constraint is measured based upon its distance from the resulting regression line. Regions having a greater weighted distance between protein-changing variants than expected based upon their CpG density (the residual from the linear regression line) are predicted to be under the greatest constraint. The resulting residuals are scaled from 0 to 100, ranked by residual (highest to lowest), and assigned a percentile such that regions with the largest residual value are assigned the highest percentile, reflecting the highest predicted constraint. Genomic positions harboring observed variants in gnomAD are assigned the lowest residual and a percentile of 0. This is based on the fact that such variants were obtained from individuals that are either healthy or did not have developmental abnormalities, and should therefore be interpreted as unconstrained loci.

### Evaluation of CCRs with ClinVar

Odds ratio comparisons were used to test the power of our CCRs to predict the pathogenicity of new variants using ClinVar variants as a truth set. Our evaluation set consisted of solely ClinVar variants that were designated as “Pathogenic” or “Likely pathogenic” for true positive variants and “Benign” as true negative variants. All variants from both sets were also required to have at least “Criteria provided, single submitter” review status or greater with no conflicts. Any variant designated as “no assertion criteria provided”, “no assertion provided”, or “no interpretation for the single variant” was excluded from the evaluation set. Variant alleles were also excluded if they matched those observed in ExAC v1 and gnomAD datasets. True positive (pathogenic) variants were also required to have a predicted impact of 'stop_gained', 'stop_lost', 'start_lost', 'initiator_codon', 'rare_amino_acid', 'missense', 'protein_altering', 'frameshift', ‘inframe_deletion’, ‘inframe_insertion’, or ‘coding_sequence_variant’ combined with either ‘splice_acceptor_variant’ or ‘splice_donor_variant’.

Odds ratios in Figure 2A were based on a curated set of genes underlying autosomal dominant disease phenotypes from Berg et al.^58^ Odds ratios for each percentile bin were calculated by, 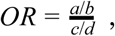 where *a* is the number of pathogenic variants in a bin, *b* is the number of benign variants in a bin, *c* is the number of pathogenics not in the bin and *d* is the number of benigns not in the bin. In other words, we are measuring the ratio of pathogenic variants in the bin to benign variants in that bin divided by the ratio of pathogenic variants not in that bin to the benign variants not in that bin. We also calculated ninety-five percent confidence intervals from the standard error, 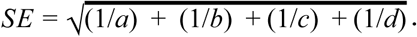 The lower confidence interval is calculated using the expression *e*^*ln*(*OR*)–1.96\**SE*^ and the upper confidence interval is calculated by *e*^*ln*(*OR*)+1.96\**SE*^.

### Evaluation of CCRs on neurodevelopmental disorder versus control *de novo*s

We used odds ratio comparisons to test the power of our CCRs to predict the pathogenicity of new variants that lie within their boundaries, and in this case, a well-curated set of de novo missense variants was used as a truth set. The set of de novo missense variants curated by Samocha et al.^42^ was used as an independent truth set for evaluating CCRs and other variant pathogenicity prediction tools. Predicted pathogenic variants in this truth set are comprised of de novo missense mutations observed in individuals with developmental delay, severe intellectual disability, and epileptic encephalopathy^44–49^. Predicted benign variants reflect de novo missense mutations from unaffected siblings of autism probands^50,51^. Mutations from both pathogenic and benign de novo sets were filtered on their presence in ExAC v1 and gnomAD. Odds ratios and confidence intervals were calculated as above.

### Comparing CCRs to missense depletion scores

We compared CCRs in the 99th percentile and higher to missense depletion scores defined by Samocha et al.^42^ by intersecting CCR regions with missense depletion regions using bedtools^59^. CCRs right of the black vertical line in Figure 4 reflect highly constrained CCRs that fall below the threshold (0.4) for significant missense depletion defined by Samocha et al.

### Coding constraint regions in Pfam domain families

Human genome build 37 genome coordinates for Pfam domains were curated from the UCSC Table Browser (Pfam Domains in UCSC Genes track). Pfam domain families were then intersected with all CCRs to measure the distribution of regional constraint across each protein domain family.

### Comparing vertebrate conservation to regional coding constraint

We investigated the relationship between constrained coding region percentiles and vertebrate conservation scores by intersecting CCRs with per-base GERP++ scores. The mean GERP++ score was calculated for each CCR. We defined CCRs in the 95th percentile or higher as constrained yet not conserved if the CCR had a mean GERP++ Rejected Substitution Score of less than 0.7 RS, as this falls 1 RS below the GERP++ confidence threshold for interspecies mammalian constraint^25^.

### Comparing CCRs and other metrics for variant prioritization

To understand how CCRs compare to other methods of variant pathogenicity prediction, we conducted a ROC curve analysis on the ClinVar truth set, and a well-curated set of de novo variants in developmental disorders (described above). The true positives were taken from both the neurodevelopmental de novo and ClinVar pathogenic and likely pathogenic variant sets respectively, filtered on matching ExAC v1 and gnomAD alleles, and the true negatives are represented by the unaffected autism sibling de novos and the ClinVar variants designated as benign.

We chose four metrics with which to compare CCRs. The first, MPC^42^ because it is the only other variant pathogenicity prediction tool that models regional constraint. Secondly, REVEL^53^ because it is a recently developed tool that performs extremely well on ClinVar compared to all other metrics. Third, GERP++^25^ as a measure of conservation for a point of comparison between constraint and conservation in human-based pathogenicity prediction. Lastly, CADD^52^ as it is a widely used variant pathogenicity prediction method.

ROC curves were calculated using scikit-learn in Python 2.7, the variants used were only protein-changing variants as defined by *pathoscore* (https://github.com/quinlan-lab/pathoscore), explained in the methods above. Fundamentally, a ROC curve takes all of the values from lowest to highest and utilizes a binary classification of true or false, depending on whether the value overlaps what is considered a true positive (in this case a pathogenic variant) or a true negative (in this case a benign variant). The scikit-learn ROC module plots the true positive rate (*TPR = TP*/(*TP + FN*)) versus the false negative rate (*FPR = FP*/(*FP + TN*)) at different thresholds determined by the machine learning algorithm.

### Gene pathway and subnetwork overrepresentation analysis

We used the “pathway-based sets” gene set overrepresentation method from ConsensusPathDB (http://cpdb.molgen.mpg.de/) to test for gene overrepresentation in distinct pathways. The ConsensusPathDB overrepresentation is calculated using a binomial test, where the null hypothesis assumes that genes in the list given are sampled from the same superset and thus the probability of observing a gene in a pathway in the given list is the same as the original superset^60^.

### Estimation of FDR and FPR

We estimate as FDR as: (*FP*/(*FP* + *TP*), where true positives (TP) are the developmental de novos that lie within a CCR above our threshold, and the false positives (FP) are the unaffected autism sibling de novos also above that threshold. Similarly, to estimate FPR (false positive rate), we create an estimate using the equation. *FPR* = *FP*/(*FP* + *TN*). We assume that, as with FDR, the false positives are the true negatives above the CCR percentile cutoff, and that the true negatives are the set of all true negatives, which, in this case is a superset of the false positives. Therefore, FPR is the true negatives above the cutoff, divided by the number of all true negatives.

## SUPPLEMENT

**Supplementary Figure 1.**
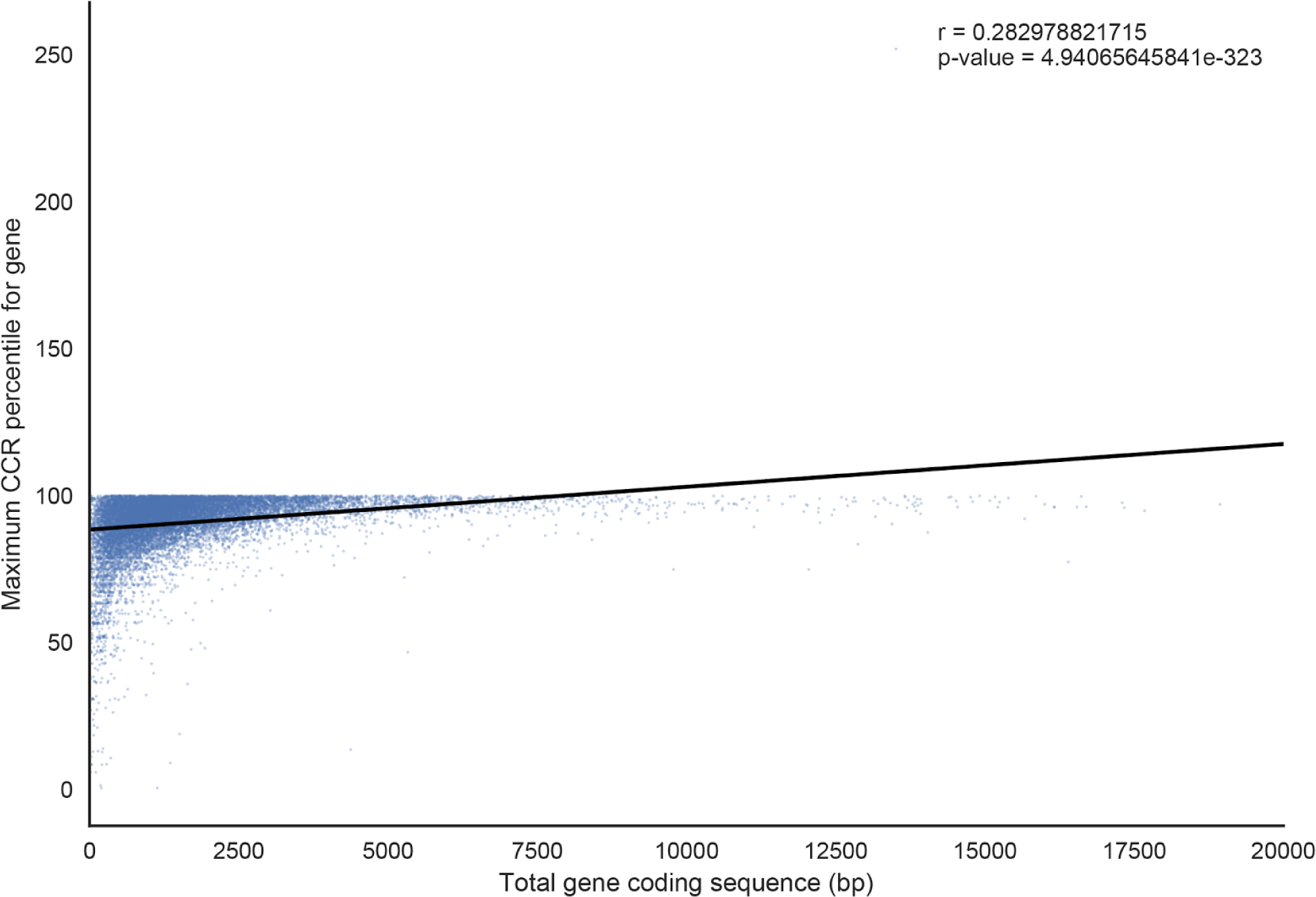
The size of each gene in total coding base pairs compared to the maximum observed CCR in the gene.

**Supplementary Figure 2.**
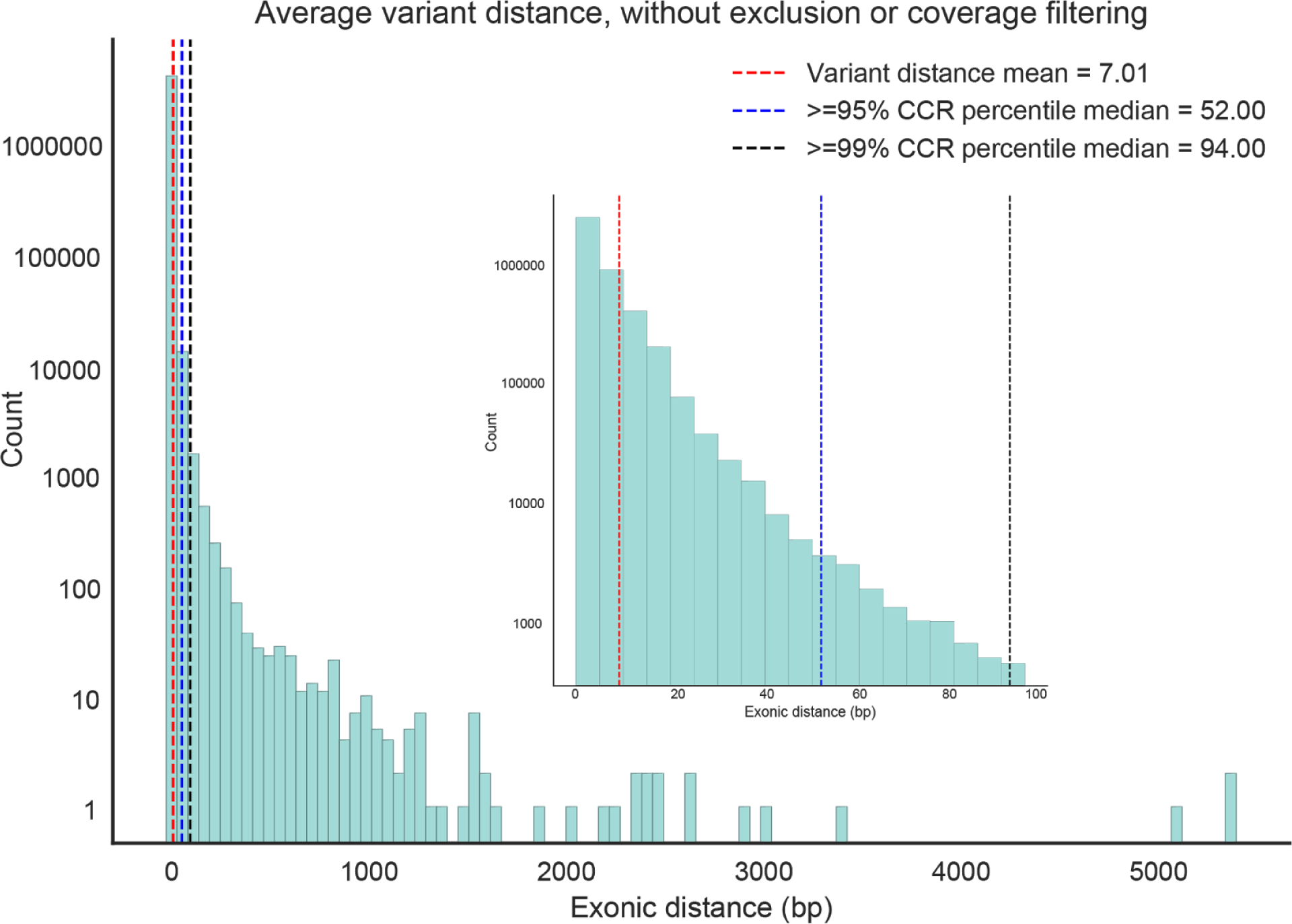
Distribution of the exonic distance between protein-changing (missense or LoF) variants in gnomAD without filtering regions by coverage, segmental duplications, or self-chains. The red dashed line is the average distance between protein-changing variants. The blue and black dashed lines represent the average length of CCRs in the 95th and 99th percentile, respectively.

